# Sporulation activated via σ^W^ protects *Bacillus* from a Tse1 peptidoglycan hydrolase T6SS effector

**DOI:** 10.1101/2022.02.23.481616

**Authors:** Alicia I. Pérez-Lorente, Carlos Molina-Santiago, Antonio de Vicente, Diego Romero

**Affiliations:** Instituto de Hortofruticultura Subtropical y Mediterránea ‘‘La Mayora’’, Universidad de Málaga-Consejo Superior de Investigaciones Científicas (IHSM-UMA-CSIC), Departamento de Microbiología, Universidad de Málaga, Bulevar Louis Pasteur 31 (Campus Universitario de Teatinos), 29071 Málaga, Spain

**Keywords:** *Bacillus*, *Pseudomonas*, sigma-factors, sporulation, T6SS, interspecies interaction

## Abstract

Bacterial communities constantly interact with various community members employing diverse offensive and defensive tools to reach coexistence. The extracellular matrix and sporulation are defensive mechanisms used by *Bacillus* cells when they interact with *Pseudomonas* strains expressing a type VI secretion system (T6SS). Here, we define Tse1 as the main toxin mobilized by the *Pseudomonas* T6SS that triggers sporulation in *Bacillus*. We characterize Tse1 as a peptidoglycan hydrolase that indirectly alters the dynamics and functionality of the *Bacillus* cell membrane. We also delineate the response of *Bacillus* cells to Tse1, which through the coordinated actions of the extracellular sigma factor σ^W^ and the cytoplasmic histidine kinases KinA and KinB, culminates in activation of the sporulation cascade. We propose that this cellular developmental response is conserved in *Bacilli* to defend against the toxicity of T6SS-mobilized Tse1 effector.

## Introduction

Multispecies microbial communities are constantly in competition for scarce resources such as space and nutrients (Granato, Meiller-Legrand and Foster, 2019). A diverse battery of offensive and defensive tools, including secretion of compounds (e.g., siderophores or antibiotics) and extracellular matrix production, are specifically deployed to modulate inter- and intraspecies coexistence (Romero, 2016; Molina-Santiago *et al*., 2019, 2021; Niehaus *et al*., 2019). An outstanding example of cellular machinery implicated in shaping microbial community structure is the type VI secretion system (T6SS), a molecular nanomachine designed to inject effectors into competitor cells (Russell, Peterson and Mougous, 2014). Initially described in *Vibrio cholerae* (Pukatzki *et al*., 2006) and *Pseudomonas aeruginosa* (Mougous *et al*., 2006), the T6SS is now believed to be present in 25% of Gram-negative bacteria. T6SSs comprise 13 core-components that form a versatile bacteriophage-like structure. The integral membrane complex is formed by TssJ, TssM and TssL, while the baseplate is formed by TssA, TssE, TssF, TssG, and TssK. The TssB and TssC subunits are responsible for assembly and disassembly of the contractile sheath, which is formed by hexamers of Hcp. The tip of the spike is formed by a trimer of VgrG, a valine-glycine repeat protein, and PAAR, a proline-alanine-alanine-arginine repeat protein that facilitates delivery of specific effectors (Shneider *et al*., 2013; Wood *et al*., 2019; Zheng, Yang and Dong, 2021). Finally, the ClpV ATPase is involved in the disassembly and recycling of T6SS components (Felisberto-Rodrigues *et al*., 2011; Silverman *et al*., 2012; Chen *et al*., 2015) (**Fig EV1A**).

T6SSs of various microbes have been reported to mobilize effectors against other Gram-negative bacteria (Jurėnas and Journet, 2021) and/or eukaryotic cells during bacterial infection (Coulthurst, 2013; Durand *et al*., 2014; Russell, Peterson and Mougous, 2014). Recently, it has been demonstrated that *Serratia marcescens* expresses a T6SS that secretes two effectors capable of acting against fungal cells (Trunk *et al*., 2018). Two additional studies have reported effects of T6SSs against Gram-positive bacteria. Among Pseudomonads, *Pseudomonas chlororaphis* PCL1606 (referred to as Pchl), a soil-dwelling bacterium known for its biocontrol activity against avocado pathogens, has recently been reported to encode a T6SS that triggers sporulation in the Gram-positive bacteria *Bacillus subtilis* upon close cell-to cell contact (Cazorla *et al*., 2006; Molina-Santiago *et al*., 2019). Furthermore, *Acinetobacter baumannii* has been shown to mobilize a bifunctional effector that seems to kill *B. subtilis* and other Gram-positive bacterium (Le *et al*., 2021).

*Bacillus subtilis* NCIB3610 (referred to as Bsub) is a Gram-positive soil dwelling bacterium frequently used as a model in studies of bacterial cell differentiation and gene expression regulation (Branda *et al*., 2006; Vlamakis *et al*., 2013; Mielich-Süss and Lopez, 2015). In ecologically relevant environments, Bsub self-defence relies on at least two complementary mechanisms: i) the production of an extracellular matrix, which protects bacterial colonies from toxic compounds produced by competitors and ii) the formation of spores in response to external cues that threaten bacterial cell viability (Molina-Santiago *et al*., 2019). Initiation of the sporulation pathway is orchestrated by the coordinated action of kinases that ultimately increase the intracellular level of the phosphorylated form of the master regulator Spo0A, which then activates a deeply understood genetic cascade (Burbulys, Trach and Hoch, 1991; Helmann, 2007; Grau *et al*., 2015). *Pseudomonas* spp. and *Bacillus* spp. are among the most studied beneficial soil microbes found in rhizosphere communities (Mendes, Garbeva and Raaijmakers, 2013; Bernal *et al*., 2017). These species can establish synergistic and antagonistic relationships; however, these relationships have been scarcely analysed and it is still necessary to study the implications of the coexistence between these two genera (Simões *et al*., 2008; Powers *et al*., 2015; Comeau *et al*., 2021; Molina-Santiago *et al*., 2021).

In a previous study, we reported that Bsub cells activate the sporulation cascade as a defence strategy against T6SS^Pchl^ when interacting with Pchl (Molina-Santiago *et al*., 2019). In this work, we identify and elucidate the mechanism by which a T6SS^Pchl^ effector, Tse1, modifies Bsub bacterial anatomy and physiology to trigger sporulation (**Fig EV1B**). We additionally show that Bsub responds to the Tse1 offensive via the coordinated activities of the extracellular sigma factor σ^W^ and the cytoplasmic histidine kinases KinA and KinB, which ultimately result in sporulation-triggering phosphorylation.

## Results

### T6SS cluster possesses several putative toxins

T6SSs have been established as an effective bacterial appendage that modulates inter- and intraspecies bacterial interactions in a spatially dependent manner. We previously demonstrated that a Pchl strain expressing a T6SS triggered Bsub sporulation; however the exact mechanism underlying this interaction remained unsolved (Molina-Santiago *et al*., 2019). An *in-silico* analysis of the Pchl genome (Calderón *et al*., 2015) revealed a complete T6SS gene cluster of 27.9 KB (T6SS^Pchl^) that is similar in structure and composition to the T6SSs of other soil dwelling bacteria, including *Pseudomonas syringae* pv. tomato DC3000, *Pseudomonas aeruginosa* PAO1, and *Pseudomonas putida* KT2440 (**Fig 1A**). A phylogenetic analysis based on the *tssB* locus positioned T6SS^Pchl^ into group 1.1, placing it very close to the HIS-I-T6SS of *P. syringae* pv. tomato DC3000, H2-T6SS of *P. aeruginosa* PAO1, and K2-T6SS of *P. putida* KT2440 (**Fig 1B**). The T6SS^Pchl^ main gene cluster comprises i) twelve structural protein components of the T6SS syringe; ii) two putative T6SS effectors based on BLASTP (Bheema Lingeswara Reddy *et al*., 2011) and the SecReT6 database (Li *et al*., 2015), one linked to a *paar* gene and the other to a *vgrG* gene; and iii) regulatory sequence for the sigma factor σ^54^, which is involved in differential regulation of T6SSs in *Pseudomonas aeruginosa*, *Salmonella enterica*, *Vibrio chlolerae*, and *Aeromonas hydrophila* (Bernard *et al*., 2011; Leung *et al*., 2011). We additionally identified six other putative T6SS effectors encoded in five orphan *vgrG* islands (**Fig 1C**).

**Figure 1.**
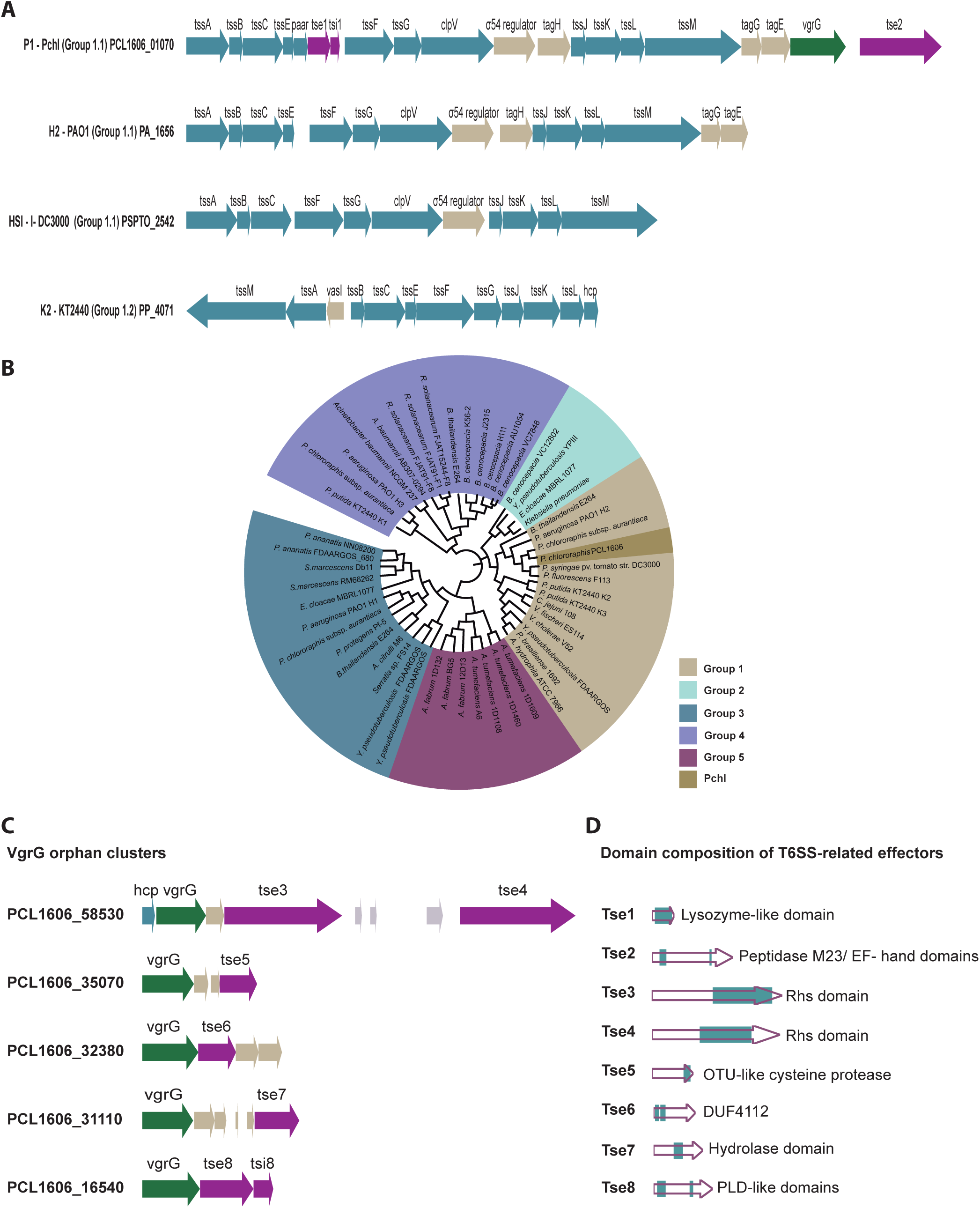
Characterization of the *P. chlororaphis* T6SS. **A)** Genetic architecture of various *Pseudomonas* group 1 T6SS clusters. Schematic representations of the main Pchl T6SS cluster, the H2-T6SS cluster of *P. aeruginosa* PAO1, the HS1-1-T6SS of *P. syringae* pv. *tomato* DC3000, and the K2-T6SS of *P. putida* KT2440. **B)** Phylogenetic distribution of T6SS clusters in five main groups based on the *tssB*. Maximum likelihood method with 1000 bootstrap replicates was built with Mega X (41) using Clustal Omega (39). The final representation was built using iTOL (40). Group 1 is represented in brown, group 2 in light blue, group 3 in dark blue, group 4 in purple, and group 5 in pink. Pchl (dark brown) belongs to group 1. **C)** Schematic representation of the VgrG orphan clusters of Pchl. **D)** Schematic representation of the predicted domains of eight putative T6SS effectors. Genes encoding structural T6SS proteins are coloured in blue, transcriptional genes in brown, adaptor genes in soft brown, VgrG genes in green, and effectors/immunity genes in pink. All arrows representing T6SS genes and the domain sizes of putative effectors are to scale.

We further analysed the domain architecture of the eight putative effectors using Pfam and Phyre2 (**Fig 1D**). Tse1 (<u>T</u>ype six <u>s</u>ecretion system <u>e</u>ffector) is encoded in the main gene cluster and is linked to a *paar* gene and followed by its putative cognate immunity protein Tsi1 (<u>T</u>ype six <u>s</u>ecretion system <u>i</u>mmunity protein) (toxin-immunity pair). Tse1 is a hypothetical protein with predicted lysozyme-like activity based on amino acid sequence homology. Tse2, also present in the main gene cluster, is a putative peptidase of the M23 family, with a C-terminal EF-hand domain. Tse3 and Tse4 are located in the same *vgrG* orphan cluster, and both contain a C-terminal Rhs (Rearrangement hot spot) domain. Effectors containing these domains usually target nucleic acids in the target cells (Alcoforado Diniz and Coulthurst, 2015; Donato *et al*., 2020). Downstream of *vgrG* is *hcp*, which encodes a structural component essential for T6SS function. Tse5 is a putative cysteine protease, likely having a function similar to that of an effector recently described in *Edwardsiella piscicida* (Li *et al*., 2021). Tse6 contains two uncharacterized DUF4112 domains in its N-terminus, similar to those primarily found in *Acinetobacter* strains. Tse7 contains a conserved hydrolase domain. Tse8 is a putative phospholipase with two PLD-like domains and is associated with the Tsi8 immunity protein. We did not identify genes encoding immunity proteins for the Tse3, Tse4, Tse5, Tse6 and Tse7 putative effectors.

### Tse1 is a T6SS effector that specifically triggers *B. subtilis* sporulation

To validate our *in-silico* analysis by verifying the functionality of T6SS^Pchl^, we studied the Bsub sporulation rate in competitive experiments with Pchl strains carrying mutations in different T6SS^Pchl^ components. As previously shown in pairwise interactions on solid medium, the Bsub sporulation rate increased up to 80% in the presence of the WT Pchl strain. The Bsub sporulation rate decreased by 40% upon interaction with a *tssA* mutant, a value statistically similar to the intrinsic sporulation rate of Bsub in the absence of competitors (**Fig EV2A**). TssA has been recently shown to be essential for the assembly and polymerization of the T6SS sheath in *P. aeruginosa* and *Escherichia coli*; thus, *tssA* mutants lack active T6SSs (Planamente *et al*., 2016; Zoued *et al*., 2016; Schneider *et al*., 2019). Accordingly, the WT Pchl strain but not the *tssA* mutant strain killed an *E. coli* biosensor strain constitutively expressing GFP (Hachani, Lossi and Filloux, 2013). *P. putida* KT2440 (Espinosa-Urgel, Salido and Ramos, 2000), which possesses two out of three T6SSs, K2-T6SS and K3-T6S, phylogenetically close to Pchl, failed to induce Bsub sporulation **(Fig EV2A)**, although it retained toxicity against *E. coli* **(Fig EV2B)**. As expected, the triple mutant, KT2440ΔT6SS, failed to kill *E. coli.* These findings support the existence of a functional T6SS in Pchl that specifically attacks Bsub cells.

The lack of activity of T6SS^KT2440^ against Bsub suggested that Bsub sporulation might be triggered by a specific T6SS-effector encoded in the Pchl genome rather than by physical damage inflicted by the penetration of the syringe. Additional evidence also supported this hypothesis: i) a strain mutant for *hcp*, which encodes an essential structural component for the assembly and contraction of the tube, failed to trigger sporulation or to kill *E. coli* and ii) a strain mutant for *paar*, which encodes a structural component of the spike of certain strains but is not essential for tube assembly, failed to trigger Bsub sporulation but still killed *E. coli* (**Fig 2A-B**). PAAR has been reported to facilitate the mobilization of specific effectors normally encoded in proximal downstream loci. Our *in-silico* analysis revealed that *tse1*, which encodes a hypothetical protein with a putative lysozyme domain, is located immediately downstream *paar* (**Fig 1A**). A *tse1* mutant failed to induce Bsub sporulation but killed *E. coli*. Based on these findings, we concluded that Tse1 is a T6SS^Pchl^-mobilized effector that is specifically involved in triggering Bsub sporulation.

**Figure 2.**
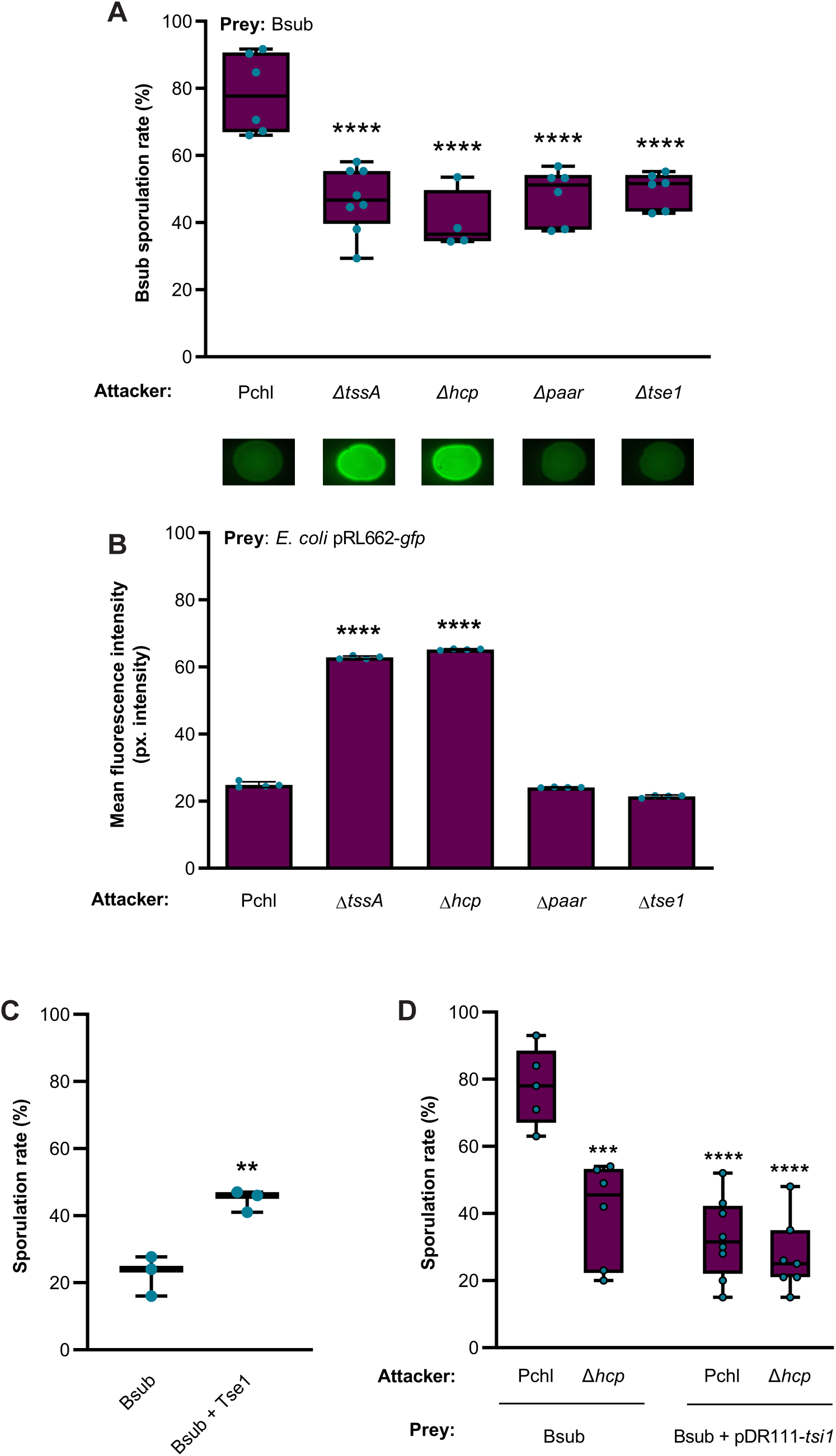
Tse1 is a toxic effector capable of triggering *Bacillus* sporulation. **A)** Competition assays. Quantification of the percentage of sporulated Bsub cells during competition with different Pchl strains as attackers (Pchl, Δ*tssA*, Δ*hcp*, Δ*paar* and Δ*tse1*). **B)** T6SS killing assays towards *E. coli* as an indicator. Fluorescence signal measurements from *E. coli* pRL662-gfp after incubation with different Pchl strains (Pchl, Δ*tssA*, Δ*hcp*, Δ*paar* and Δ*tse1*). Images show a representative spot of each assay after 24 h of incubation. **C)** Quantification of the percentage of sporulated Bsub cells after treatment with purified Tse1 for 3 h. **D)** Competition assays of Pchl (functional T6SS) or Δ*hcp* (non-functional T6SS) against the preys *Bacillus* (WT) or Bsub pDR111-tsi1 expressing the putative cognate immunity protein Tsi under the control of a constitutive promoter. In all represented competition assays (A, B, D), the attacker and prey strains were cultured at a ratio of 1:1. After 24 h, *Bacillus* cells were serially diluted and placed into LB for estimation of cell density (CFU) sporulation percentages. In all experiments, at least three biological replicates are shown, and the error bars represent SD. Statistical significance was assessed via t-tests and one-way ANOVA tests. ** p value < 0.01, *** p value < 0.001, **** p value <0.0001.

To explore Tse1 function, we first demonstrated its bacteriolytic activity in bacterial intoxication assays with *E. coli* strains harbouring a *tse1^+^*-expressing plasmid (pDEST17-*tse1*) or a plasmid encoding *tse1* lacking its N-terminus (pDEST17-C*tse1*) controlled by an inducible promoter. After induction with L-arabinose, decreased cell viability in *E. coli* harbouring pDEST17-*tse1* resulted in lower cell density compared with that of *E. coli* harbouring pDEST17-C*tse1* (**Fig EV2C**). Inspired by this finding, we then purified Tse1 to homogeneity and evaluated its activity against Bsub (**Fig EV2D**). A 7 µM solution of purified Tse1 significantly increased the Bsub sporulation rate (43%) compared with that of untreated cells (**Fig 2C**). T6SS effectors are usually encoded near their cognate immunity proteins, and both are concurrently produced to avoid self-intoxication (Russell *et al*., 2011). *tsi1* is located immediately downstream *tse1* (**Fig 1A**); thus, we speculated that these proteins might form an effector-immunity (EI) pair. No significant differences in the sporulation rate of Bsub cells overexpressing the EI protein Tsi1 (Bsub + pDEST17*-tsi*) were observed in competition assays with WT Pchl (functional T6SS) or a Δ*hcp* (non-functional T6SS) strain, indicating that Tsi1 produced in Bsub cells can neutralise the toxic effect of Pchl-secreted Tse1 (**Fig 2D**).

### Tse1 is a hydrolase that degrades *B. subtilis* peptidoglycan

We speculated that the cell wall is the main target of Tse1 for two reasons: i) the predicted lysozyme domain in Tse1 and ii) the extracellular toxicity of Tse1 to Bsub cells. Transmission electron microscopy of thin sections of Bsub cells after incubation with Tse1 for 3 h revealed abundant cell ghosts (**Fig 3A**). Complementary transmission electron microscopy confirmed the cell wall defects in Bsub cells (**Fig EV3**). Immunofluorescence assays showed that His-tagged Tse1 colocalised with Alexa Fluor 647-conjugated wheat germ agglutinin (WGA), a fiducial marker of local cell wall growth that binds *N*-acetylglucosamine residues at the septa and poles of Bsub cells (**Fig 3B** **and Fig EV4A-B)** (Ursell *et al*., 2014; El Biari *et al*., 2019; Yaghoubi *et al*., 2020).

**Figure 3.**
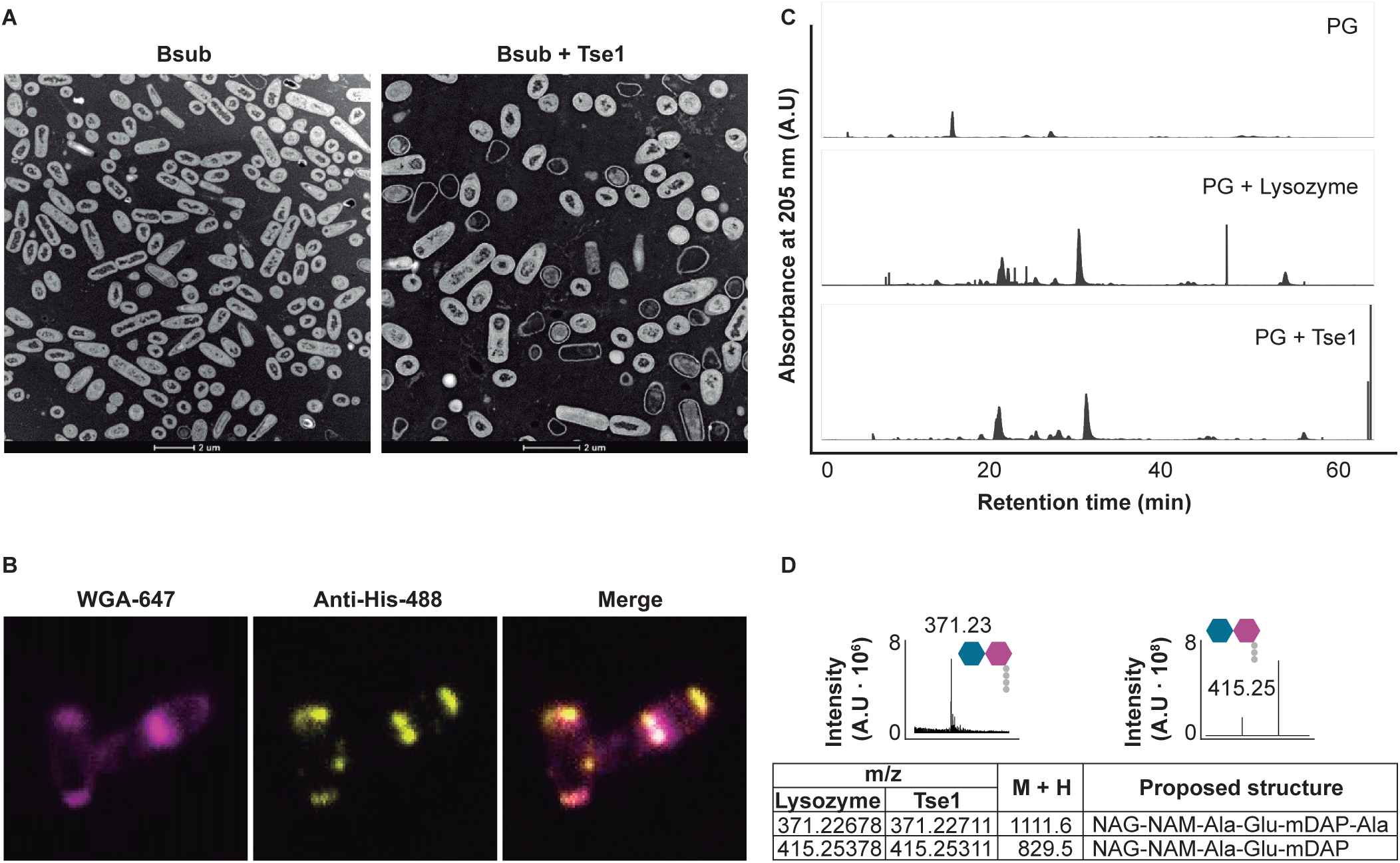
Tse1 hydrolyses *Bacillus* peptidoglycan. **A)** Transmission electron micrograph of positively stained thin sections of *Bacillus* cells showing an overrepresentation of ghost cells after Tse1 treatment (Bsub + Tse1) compared with untreated cells (Bsub, left panel). Scale bars equal 2 μm. **B)** Immunofluorescence assay of a Tse1-treated *Bacillus* bacterium immunolabeled with anti-His antibody conjugated to Alexa Fluor 488 (Anti-His-488, yellow channel) and stained with WGA conjugated with Alexa Fluor 647 (WGA-647, pink channel). Protein accumulation is visible at the cell septa and poles. Images of untreated *Bacillus* cells and colocalization analysis are shown in Expanded view Figures 4A and 4B. **C)** HPLC chromatograms of reduced soluble *Bacillus* peptidoglycan products obtained from digestion with lysozyme (PG + lysozyme, positive control), with Tse1 (PG + Tse1), or treated with buffer (PG, negative control). **D)** Partial HPLC chromatograms of muropeptides eluted at 55.89 min (371.23 m/z) and 63.65 min (415.25 m/z) obtained after Tse1 treatment. Peak assignments were made based on MS data and literature review. Predicted structures are shown with hexagons (NAM and NAG) and circles. Table showing the most abundant and relevant peaks found in samples treated with lysozyme or Tse1, their mass in Daltons (M + H), and the proposed structures. NAG, *N*-acetylglucosamine; NAM, *N*-acetylmuramic acid; Ala, alanine; Glu, glutamic acid; mDAP, *meso*-diaminopimelic acid; A.U., arbitrary units.

To further validate the hypothetical enzymatic activity of Tse1, purified Bsub sacculi were treated with Tse1 and the products were analysed via LC-MS. Consistent with the suspected putative enzymatic activity, the elution profile of muropeptides was similar to that obtained when lysozyme was used as a control (**Fig 3C**). In addition, two specific peaks were relatively more abundant: 371.23 Da, corresponding to the disaccharide tetrapeptide NAG-NAM-Ala-Glu-mDAP-Ala and 415.25 Da, corresponding to the disaccharide tripeptide NAG-NAM-Ala-Glu-mDAP (**Fig 3D** **and Fig EV5)** (Atrih *et al*., 1999; Bacher *et al*., 2001; Rodríguez-Rubio *et al*., 2016). Altogether, these findings confirmed the peptidoglycan hydrolase activity of Tse1, which might be a member of the muramidase enzyme family.

The damage inflicted to the peptidoglycan led us to analyse the integrity and functionality of the cell membrane. First, staining with propidium iodide (PI), a DNA intercalator used to evaluate cell membrane integrity (Löpez-Amorós *et al*., 1997), revealed a significant increase in the number of permeable Bsub cells after treatment with purified Tse1 (**Fig 4A**). Second, staining with TMRM showed a reduction of the mean fluorescence intensity, indicating a loss of membrane potential (**Fig 4B** **and Fig EV6**). Third, staining with DilC12 revealed the presence of highly fluorescent foci (increased fluidity), indicating that Tse1 increased the fluidity of the cell membrane (**Fig 4C**).

**Figure 4.**
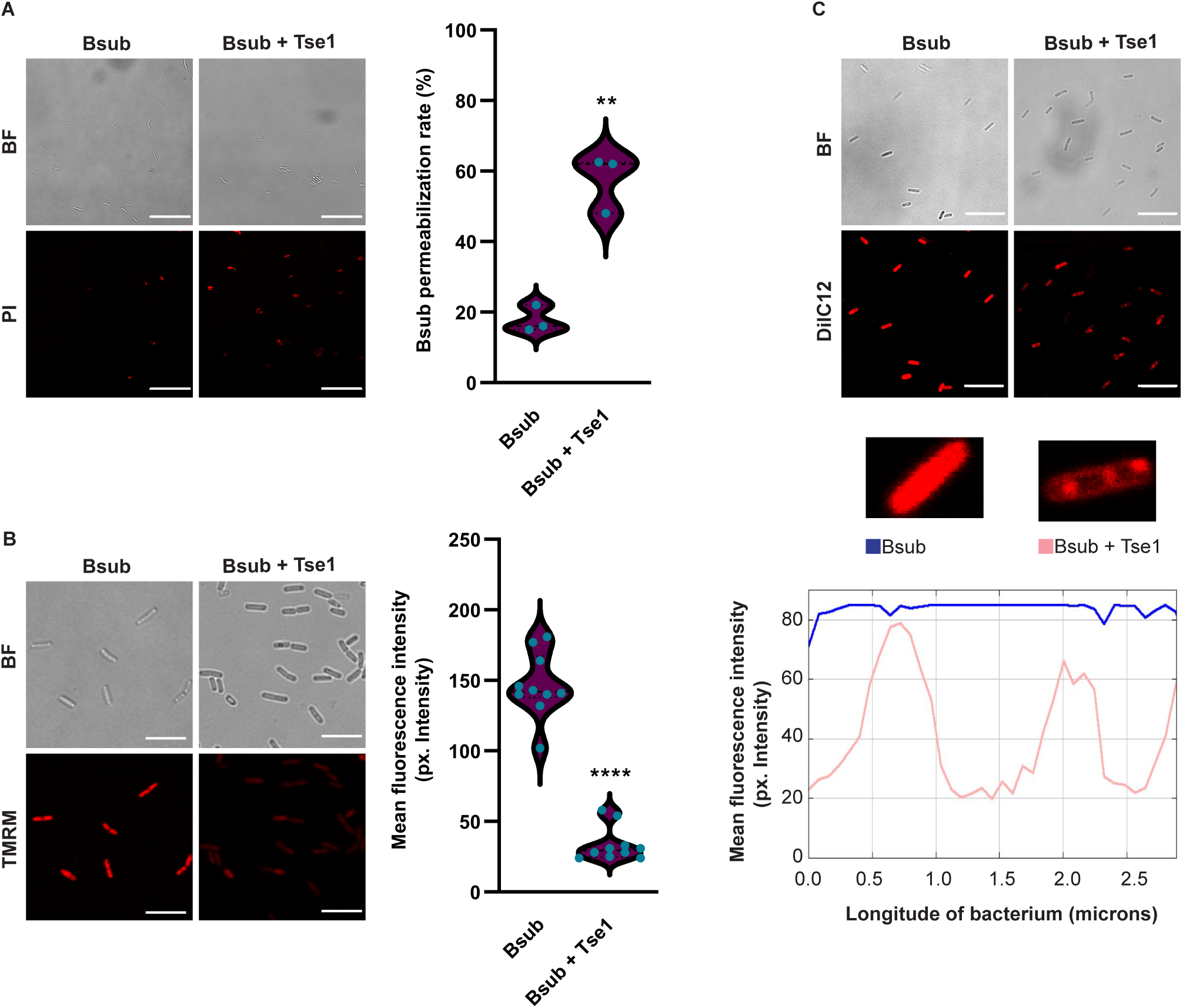
Tse1 affects *Bacillus* membrane integrity. **(A)** CLSM of propidium iodide-stained Bsub cells revealed increased membrane permeability in Tse1-treated *Bacillus* cells (right panel) compared with untreated cells (left panel) and graph. Scale bars equal 20 μm**. B)** Quantification of the fluorescence signal obtained after TMRM staining and CLSM showed a significant decrease in membrane potential in Tse1-treated Bsub cells (right panels) compared with that in untreated cells (left panels). Scale bars equal 7 μm. **C)** Staining with Dilc12 and CLSM revealed higher membrane fluidity in Tse1-treated Bsub cells (right panels) compared with untreated (left panels). Representation of the plot profile of DilC12 in untreated Bsub or Tse1-treated cells showing the Dilc12 distribution along a bacterium. The blue line represents the mean fluorescence intensity at each point of a single Bsub cell and the pink line represents the mean fluorescence intensity along a single Tse1-treated Bsub bacteria. Scale bars equal 10 μm. Experiments were repeated at least three times with similar results. Statistical significance in the TMRM and PI experiments was assessed via t-tests. **p value < 0.01, ****p value < 0.001.

### σ^W^ activates *Bacillus* sporulation in response to Tse1-inflected peptidoglycan damage

To identify the genetic pathways connecting the anatomical and functional alterations of the Bsub cell envelope with sporulation, the whole transcriptome of Bsub cells was sequenced after treatment with Tse1 for 3 h. A total of 97 genes were differentially expressed in Tse1-treated cells compared with untreated cells (**Dataset EV1 and EV2)**. Consistent with the specific cytological modifications of Bsub cells (**Fig 3A**), 58% of the differentially upregulated genes belonged to the σ^W^ regulon, which is associated with cell wall stress, cell membrane homeostasis, and resistance to cell wall-directed antibiotics (Cao *et al*., 2002; Luo and Helmann, 2009; Helmann, 2016; Asai, 2017). Among the genes under positive transcriptional control by σ^W^, we observed upregulation of *sasA*, which encodes the synthetase of the small alarmone guanosine tetraphosphate (ppGpp). ppGpp inhibits GMP kinase; thus, increased ppGpp leads to a decrease in the GTP pool and a corresponding increase in ATP levels (Kriel *et al*., 2013; Fung *et al*., 2020; Mediates *et al*., 2020). Among the five described kinases, KinA and KinB are known to sense variations in cellular ppGpp levels and activate Spo0F phosphorylation, leading to a large pool of phosphorylated Spo0A, which then ultimately controls the choice between two cell fates: sporulation and biofilm formation (**Fig 5A**). Strains mutant for *kinA* or *kinB*, but not for the remaining three canonical kinases, were blind to the presence of WT Pchl, and accordingly, their sporulation rates were unchanged during competition with Pchl codifying a functional T6SS (**Fig EV7A**).

**Figure 5.**
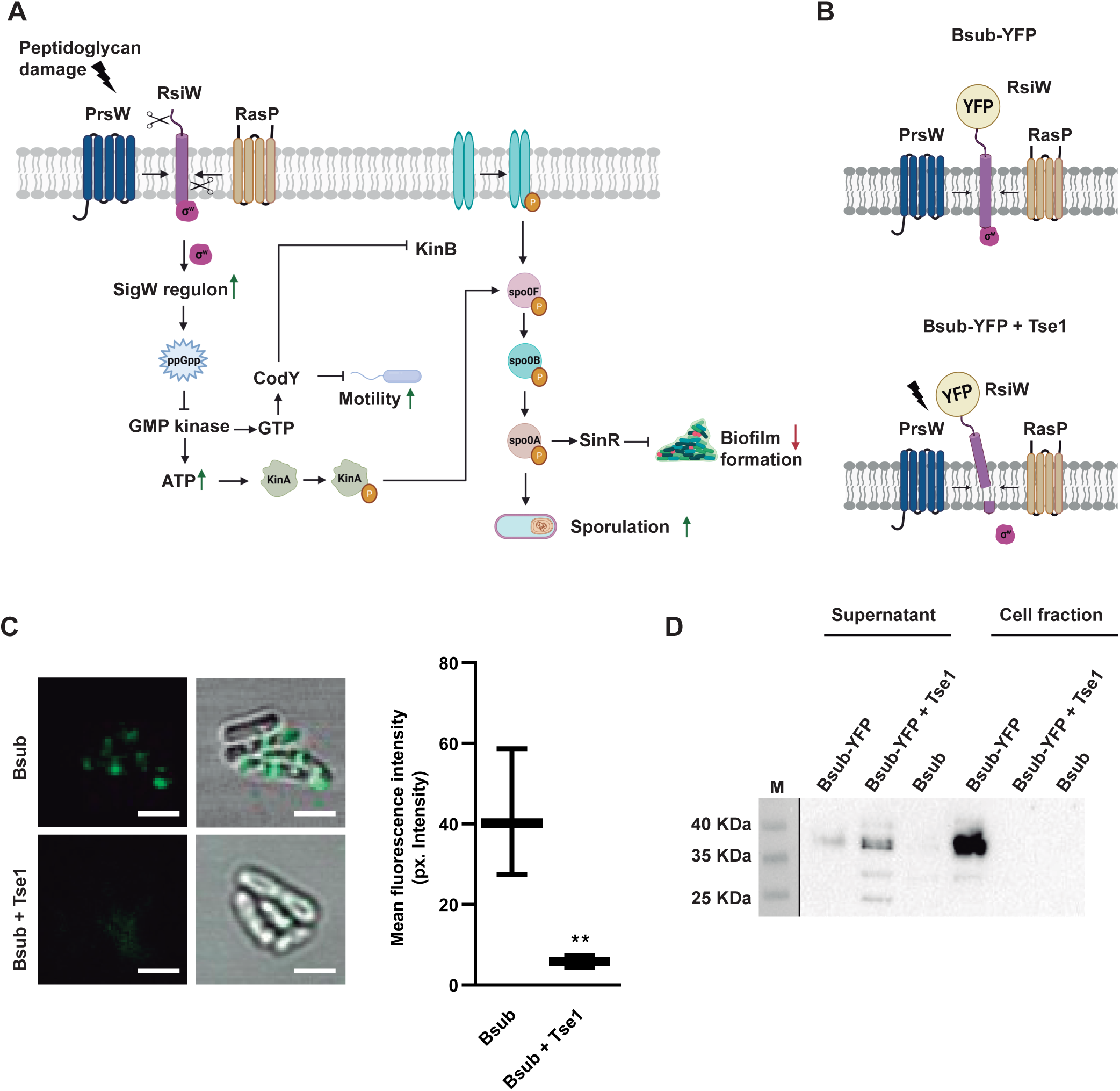
*Bacillus* sporulation is mediated by σ^W^ and the KinA and KinB kinases. **A)** Transcriptomic analysis revealed how Tse1-induced cell wall damage to Bsub activates the σ^W^ regulon, which eventually triggers the KinA-and KinB-mediated phosphorelay pathway. Based on RNA-seq data, the green and red arrows show upregulated and downregulated processes, respectively. **B)** Schematic representation of RsiW-YFP translational fusion in control Bsub cells (top) or Tse1-treated Bsub cells (bottom). **C)** Immunofluorescence assay of *Bacillus* cells expressing RsiW-YFP translational fusion using anti-YFP antibody conjugated to Alexa Fluor 488 (green channel) showed accumulation of signal in foci of untreated cells (top) and less signal intensity in Tse1-treated *Bacillus* cells (bottom). Scale bars equal 2 µm. **D)** Western blot using anti-YFP antibodies revealed an immunoreactive band of the expected size of the RsiW-YFP translational fusion in the cell fraction of untreated Bsub cells and mostly in the supernatant of Tse1-treated Bsub cells. Bsub, *Bacillus* WT; Bsub-YFP, *Bacillus* harbouring the translational fusion RsiW-YFP. Gel images have been cropped and spliced for illustrative purposes. Experiments have been repeated at least three times with similar results. Statistical significance was assessed via t-tests. **p value < 0.01.

Activation of the σ^W^ regulon relies on the alleviation of the repression imposed by RsiW, the cognate anti-sigma factor of σ^W^. RsiW is a transmembrane protein that under cell envelope stress is degraded by the PrsW and RasP proteases. Thus, we proposed that the damage inflicted to the cell wall by Tse1 might provoke RsiW degradation, resulting in disengagement of σ^W^ and activation of the genes under its positive control (**Fig. 5B**). Confocal microscopy of cells expressing YFP-tagged RsiW (RsiW-YFP) showed signal accumulation in foci on the cell membranes of untreated cells (**Fig 5B**, top and **Fig. 5C**). Consistent with enzymatic processing of the RsiW C-terminus, Tse1 treatment led to a significant decrease in fluorescent signal intensity, most likely due to the release of the C-terminally fused YFP tag (**Fig 5B**, bottom and **Fig. 5C**). In agreement with these observations, an immunoblot analysis revealed a band with an estimated molecular weight consistent with that of the RsiW-YFP translational fusion protein that was also immunoreactive to YFP antibodies in the Bsub cell fraction (**Fig 5D** **and Fig EV7B**). Tse1 treatment eliminated this signal from the cell fraction, and a band with a molecular weight corresponding to YFP monomer and immunoreactive to YFP antibodies appeared in the cell-free supernatant (**Fig 5D** **and Fig EV7B**).

## Discussion

Until very recently, T6SSs were known to antagonize certain types of eukaryotic cells (e.g., mammalian and amoeboid) and Gram-negative bacteria, while they had not been linked to Gram-positive bacteria or fungal cells (Pukatzki *et al*., 2006; Ma *et al*., 2009; Russell *et al*., 2014; Sana, Lugo and Monack, 2017). However, recent findings have broadened the roles of T6SSs as it is now known that the *Serratia marcescens* T6SS acts against fungal cells (Trunk *et al*., 2018) and that the *P. clororaphis* and *A. baumanii* T6SSs act against Gram-positive bacteria (Molina-Santiago *et al*., 2019; Le *et al*., 2021). In our previous study, we introduced the idea that the Pchl T6SS activated sporulation in Bsub cells lacking a functional extracellular matrix. In this work, we have now identified the T6SS^Pchl^ effector responsible for this activity and dissected the cellular and molecular machinery that activate sporulation in response to the damage inflicted by this effector.

The newly identified T6SS^Pchl^-associated effector, Tse1, partially hydrolyses peptidoglycan, an offense that is rapidly sensed by Bsub cells, which then respond by activating sporulation as a last resort defensive tactic. Our data indicate that Tse1 is a muramidase that hydrolyses the peptidoglycan backbone, similar to Tse3 from *P. aeruginosa* and Tge2 from *P. fluorescens* (Russell *et al*., 2011; Whitney *et al*., 2013; Russell, Peterson and Mougous, 2014). In addition to its direct activity on peptidoglycan, Tse1 also induces changes in cell membrane functionality. In particular, our *in vitro* experiments show that Tse1 compromises membrane integrity, polarization, and permeability. Such effects are not unprecedented given that three other T6SS effectors have also been found to induce similar membrane alterations: VasX, Tse4, and Ssp6 of *Vibrio cholerae*, *P. aeruginosa,* and *Serratia marcescens*, respectively (Miyata *et al*., 2013; LaCourse *et al*., 2018; Mariano *et al*., 2019).

Our findings provide new evidence supporting the versatility and variability of T6SS activity, depending especially on the strain and likely the specific competitors that cohabit the same niche. In addition to our findings related to the effector Tse1, which appears to be restricted to *Pseudomonas* species, we found that a strain lacking *paar* (Δ*paar*) had an active T6SS that triggered sporulation comparably to Δ*hcp,* Δ*tssA,* and Δ*tse1* strains (**Fig 2A-B**). This observation is in contrast to what has been reported in *Acinetobacter baylyi* ADP1, but similar to findings from other species, including *Vibrio cholerae* and *Aeromonas dhakensis* (Shneider *et al*., 2013; Liang *et al*., 2021), where PAAR is not essential for T6SS activity. Given that *paar* deletion apparently abolishes Tse1 secretion, we suggest that Tse1 is a PAAR-associated effector that requires a PAAR repeat domain protein to be targeted for secretion, thereby increasing *Bacillus* sporulation during contact with *Pseudomonas* cells (Hachani *et al*., 2014; Whitney *et al*., 2014; Cianfanelli, Monlezun and Coulthurst, 2016).

Finally, we have elucidated the molecular mechanism by which Tse1 triggers sporulation. Based on our data, we propose a link between σ^W^ activation and Bsub sporulation. σ^W^ belongs to the extracytoplasmic function (ECF) family of σ factors. ECF σ factors are two-component regulatory systems that allow bacteria to respond to extracellular conditions that disrupt cell wall homeostasis, e.g., antibiotic exposure, heat shock, or salt stress. Among the seven ECFs present in *Bacillus subtilis* (σ^M^, σ^V^, σ^W^, σ^X^, σ^Y^, σ^Z^, and σ^YlaC^), σ^YlaC^ has been directly linked to sporulation under oxidative stress (Helmann, 2016; Emily C. Woodsa and Shonna M. McBride, 2017; Kwak *et al*., 2018). We describe how Tse1-induced peptidoglycan damage leads to σ^W^ induction and activation of the sporulation cascade, thereby connecting extracellular damage sensing by RsiW with sporulation pathway activation. Based on our results, we conclude that T6SSs, and specifically Tse1, might play roles in microbial competition by modulating population dynamics.

We propose that the mechanism of sporulation activation described in this study is a defence strategy conserved across *Bacilli*. Supporting our model, *B. cereus* AH187 (BcerAH) showed increased sporulation in the presence of Pchl with a functional T6SS (**Fig EV8A**) with no difference in cell density (**Fig EV8B-C**). Interestingly, a significant decrease in cell density was observed in competition experiments with the asporogenic strain *B. cereus* DSM 2302 (Antequera-Gómez *et al*., 2021) (BcerDSM) (**Fig EV8D**). Thus, our data suggest that sporulation is a non-immunity defence mechanism against T6SS-expressing competitors with Tse1-like with effectors. This developmental process has been extensively documented due to its extreme importance in agriculture and biotechnology, and it is a striking feature of members of the *Bacillus* genera (Schultz *et al*., 2009).

## Methods

### Bacterial strains and culture conditions

A complete list of the bacterial strains used in this study is shown in **Table EV1**. Bacterial cultures were grown in liquid LB (Lysogeny Broth/Luria-Bertani: 1% tryptone, 0.5% yeast extract and 0.5% NaCl) medium at 30 °C (*Pseudomonas* and *Bacillus*) or 37 °C (*E. coli*) with shaking on an orbital platform. The pH was adjusted to 7 prior to sterilization. When necessary, antibiotics were added to the media at appropriate concentrations.

### Construction of *P. chlororaphis* T6SS mutants

Chromosomal deletions were performed using the I-SceI methodology, based on recombination between free-ended homologous DNA sequences (Martínez-García and de Lorenzo, 2012; Molina-Santiago *et al*., 2016). Upstream and downstream segments of homologous DNA were separately amplified and then joined to a previously digested pEMG vector using Gibson Assembly Master Mix (Gibson *et al*., 2009). The oligonucleotides used are shown in **Table EV2**. The resulting plasmids were then electroporated into Pchl. After selection for positive clones, the pSEVA628S I-SceI expression plasmid was also electroporated, and kanamycin-sensitive clones were PCR analysed to verify the deletions. The pSEVA628S plasmid was cured by growth without selective pressure and its loss was confirmed by sensitivity to 60 µg/mL gentamicin and colony PCR screening.

### Construction of *B. subtilis* mutants

All of the primers used to generate the different strains are listed **Table EV2**. To build the Bsub-pDR111+tsi1 strain, *tsi* was amplified and cloned into the pDR111 overexpression vector using Gibson Assembly Master Mix. The resulting plasmid was transformed by natural competence into *Bsub* 168 replacing the *amyE* neutral locus. Transformants were selected via spectinomycin resistance. To build the Bsub pDR183 + RsiW-YFP strain, RsiW and YFP were amplified by PCR and cloned into pUC19. The resulting plasmid was cloned into pDR183 by transforming *B. subtilis* 168 via its natural competence and then using positive clones as donors for transferring the constructs into Bsub via SPP1 phage transduction, replacing the *lacA* neutral locus (Yasbin and Young, 1974).

### Bacterial competition and intoxication assays

*Bacillus* and *Pseudomonas* strains were grown overnight in 5 mL LB before normalization to an OD_600_ = 3.0 in 1 mL sterile distilled water. Attacker and prey strains were mixed at a 1:1 ratio, and 2.5 µL competition drops were spotted onto LB plates and incubated at 28 °C for 24 h. The resulting colonies were resuspended in 1 mL of sterile distilled water and serially diluted and plated on LB. Next, *Bacillus* cells were selected by temperature at 40 °C, and CFUs were enumerated after 24 h. *Pseudomonas* T6SS killing assays were performed using *E. coli* DH5α pRL662-gfp as prey, whose plasmid confers strong constitutive GFP expression.

Overnight LB cultures *of E. coli* BL21 (AI) harbouring pDEST17 encoding Tse1 were adjusted to OD_600_ = 0.1 and incubated at 37 °C until OD_600_ = 0.6. Next, *tse1* expression was induced with 0.2 % L-arabinose.

### SDS-PAGE and immunodetection

SDS-PAGE gels were routinely used to analyse protein samples. Precipitated proteins were resuspended in 1x Laemmli sample buffer (BioRad) and heated at 100 °C for 10 min. Proteins were separated via SDS-PAGE in 15% acrylamide gels and then transferred onto polyvinylidene difluoride (PVDF) membranes using the Trans-Blot Turbo Transfer System (BioRad) and PVDF transfer packs (BioRad). For immunodetection of recombinant His-tagged Tse1, the membranes were probed with anti-His antibodies (rabbit) used at a 1:1000 dilution in Pierce Protein-Free (TBS) blocking buffer. A secondary anti-rabbit IgG antibody conjugated to horseradish peroxidase (BioRad) was used at a 1:3000 dilution in the same buffer. The membranes were developed using the Pierce ECL Western Blotting Substrate (ThermoFisher). For immunodetection of YFP fused to RsiW, anti-YFP antibodies were used at a 1:1000 dilution in blocking buffer, and a secondary anti-rabbit IgG antibody conjugated to horseradish peroxidase was used at a 1:3000 dilution in the same buffer.

### Bioinformatic analysis

*Pseudomonas* sequences were obtained from NCBI. BLASTP analyses were performed at the pseudomonas.com website. The Protein Homology Recognition Engine server (Phyre2) was used to predict structure-based homology. Protein domain predictions were performed via Pfam. The phylogenetic tree was constructed using MEGAX software and the iTOL website.

All *tssB* genes from Gram-negative bacteria with characterized T6SSs were collected from the SecReT6 database. Based on the Hcp hidden Markov model (HMM) Cluster representatives were aligned using Clustal Omega (Madeira *et al*., 2019), and a phylogenetic tree was constructed using MegaX. The phylogenetic tree was visualized using iTOL (Letunic and Bork, 2007).

### Protein expression and purification

To express and purify recombinant His_6_-tagged Tse1, *E. coli* BL21 (AI) was transformed with pDEST-tse1. Freshly transformed *E. coli* BL21 (AI) colonies harbouring pDEST17-tse1 were grown in LB at 37 °C overnight and reinoculated at a ratio of 1:100 in fresh LB for 2–3 h prior to induction of gene expression by addition of 0.2 % L-arabinose and growth for 2 h at 28 °C. Cells were collected via centrifugation (7000 x *g*, 10 min, 4 °C) and resuspended in buffer A (Tris 50 mM, 150mM NaCl, pH8) supplemented with 0.5 mg/mL lysozyme, 5 mM PMSF, and 10x Cell Lysis Reagent (Sigma), and incubated for 1 h at 37 °C. Next, the cells were disrupted via sonication on ice (4 x 60 s, 80% amplitude) and passed through a 0.45-μm filter prior to protein purification via affinity chromatography using an AKTA Start FPLC system (GE Healthcare). The lysate was loaded into a HisTrap HP 5 mL column (GE Healthcare) previously equilibrated with binding buffer (50 mM Tris, 0.5 M NaCl, 50 mM imidazole, pH 8). Protein was eluted with elution buffer (50 mM Tris, 0.5 M NaCl, 500 mM imidazole, pH 8). Next, the purified protein was loaded into a HiPrep 26/10 desalting column (GE Healthcare), and the buffer was exchanged to Tris 20 mM, NaCl 50 mM at pH 7. Finally, to obtain highly pure desalted protein via size exclusion chromatography, the protein was loaded into a HiPrep 16/60 Sephacryl S-300 HR column (GE Healthcare).

### Membrane fluidity assays

Membrane fluidity was evaluated using the DiIC12 reagent (ThermoFisher). To evaluate regions of increased fluidity, *Bacillus* strains grown overnight on LB plates were resuspended in 1 mL of sterile distilled water followed by addition of 1 μg/mL DiIC12. 0.5% Benzyl-alcohol was added to positive control samples. Cells were incubated at 28 °C for 3 h and were washed six times with sterile distilled water. Images were obtained using a Leica SP5 confocal microscope with a 63x NA 1.3 Plan APO oil-immersion objective and acquisition with excitation at 561 nm and emission detection between 576 and 640 nm. Image processing was performed using FIJI/ImageJ software (Schindelin *et al*., 2012). For each experiment, the laser settings, scan speed, HyD detector gain, and pinhole aperture were kept constant across all acquired images.

### Membrane potential assays

Membrane potential was evaluated using the image-iT TMRM (tetramethylrhodamine, methyl ester) reagent (Invitrogen) following the manufacturer’s instructions. Colonies grown at 28 °C on LB plates were isolated at 24 h and resuspended in sterile distilled water. TMRM reagent was added to the bacterial suspensions at a final concentration of 100 nM, and the mixtures were incubated at 37 °C for 30 min. After incubation, the cells were immediately visualized via confocal laser scanning microscopy (CLSM) with excitation at 561 nm and emission detection between 576 and 683 nm. Image processing was performed using FIJI/ImageJ software (Schindelin *et al*., 2012).

### Membrane permeability assays

Membrane-level cell damage was assessed via propidium iodide (PI) staining (Invitrogen). Colonies grown at 28 °C on LB plates were isolated after 24 h and resuspended in sterile distilled water. Cells were collected via centrifugation and incubated with 5 µg/mL PI in the dark for 20 min. Next, fluorescence was measured with excitation at 535 nm and emission detection between 560 and 617 nm. Image processing was performed using FIJI/ImageJ software (Schindelin *et al*., 2012).

### Transmission Electron microscopy

Bacterial suspensions were fixed in 2% paraformaldehyde and 2.5% glutaraldehyde overnight at 4 °C. After three washes in fixation-mix, samples were post-fixed with 1% osmium tetroxide solution for 90 min at room temperature, followed by two washes, and 15 min of stepwise dehydration in an ethanol series (30%, 50%, 70%, 90%, and 100% twice). Between the 50% and 70% steps, samples were incubated in-bloc in 2% uranyl acetate solution in 50% ethanol at 4 °C, overnight. Following dehydration, the samples were gradually embedded in low-viscosity Spurr’s resin: resin:ethanol, 1:1, 4 h; resin:ethanol, 3:1, 4 h; and pure resin, overnight. The sample blocks were embedded in capsule molds containing pure resin for 72 h at 70 °C. The samples were left to dry and were visualized under FEI TALOS F200X.

### PG digestion assays

*Bacillus* sacculi isolation was performed according to previously described methods with some modifications (Alvarez *et al*., 2016; Porfírio, Carlson and Azadi, 2019). Briefly, the bacterial pellet recovered from a 50 mL exponential phase culture was resuspended in boiling 5% SDS and incubated for 1 h. The SDS was removed via ultracentrifugation (110000 x *g*, 10 min) and repeated water washes. Next, the *Bacillus* sacculi were sonicated (4 x 90 s, 80% amplitude) and incubated with 100 mM Tris-HCl pH 7, 10 mg/mL RNase A, 10 mg/mL DNase, 1M MgSO_4_, and 100 mM CaCl_2_ for 16 h at 37 °C. To inactivate enzymes, the sacculi were treated with 5% SDS, which was then removed via ultracentrifugation (110000 x *g*, 10 min) and repeated water washes. Next, the SDS-free pellet was resuspended in 8 M LiCl and incubated for 10 min at 37 °C. The pellet was then resuspended in 100 mM EDTA and incubated for 10 min at 37 °C. To remove EDTA, samples were washed with water. After ultracentrifugation, the pellet was washed with acetone. To obtain soluble muropeptides, sacculi samples were treated with lysozyme or purified Tse1 and incubated for 16 h at 37 °C. A sacculi sample incubated with buffer instead of lysozyme or Tse1 served as a control. Finally, the pH of the soluble fraction was adjusted to 8.5–9.0 with borate buffer, and the muropeptides were reduced with freshly prepared 2 M NaBH_4_ for 30 min. Before LC-MS analysis, the pH was adjusted to 2.0 with 25% orthophosphoric acid.

### LC-MS analysis

Reduced muropeptides were analysed as previously described (Anderson *et al*., 2020), with some modifications. Briefly, sacculi samples were analysed using an UHPLC Easy nLC 1200 attached to a Q Exactive HF-X Quadrupole-orbitrap mass spectrometer (Thermo Scientific). Data were acquired using Tune 2.9 and Xcalibur 4.1.31.9. Separation was performed using an autosampler, injecting 2 µL in an analytical column (10 cm, PepMap RSLC C18, 2 µm, 100 A, 50 µm x 15 cm, Thermo Scientific). For chromatographic separation, solvent A (0.1% formic acid) and solvent B (acetonitrile 80% with 0.1% formic acid) were prepared. Muropeptides were separated using 5% B for 5 min, then increasing to 20% B, further increasing to 32% and finally increasing to 95% B over 10 min, with a constant caudal of 0.3 mL/min. Muropeptide mass spectrometry detection was performed in positive mode with the electrospray capillary voltage at 1.5 kV at 250 °C. The scan range was set to 300–2000 m/z, and the normalized fragmentation collision energies were se7 at 15, 20, and 30 eV. Data were analysed using Xcalibur Qual Browser 4.2 (Thermo Fisher).

### RNA isolation and sequencing

*Bacillus* cells were centrifuged at 4 °C and then placed at -80 °C for at least 30 min. For cell disruption, the cells were resuspended in BirnBoim A (20% sucrose, 10 mM Tris-HCl pH 8, 10 mM EDTA and 50 mM NaCl), and lysozyme (10 mg/mL) was added followed by incubation for 30 min at 37 °C. After disruption, the suspensions were centrifuged, and the pellets were resuspended in Trizol reagent (Invitrogen). Total RNA extraction was then performed as indicated by the manufacturer. DNA removal was carried out by treatment with Nucleo-Spin RNA Plant (Macherey–Nagel). The integrity and quality of the total RNA was assessed with an Agilent 2100 Bioanalyzer (Agilent Technologies) and via electrophoresis. Removal of rRNA was performed using the RiboZero rRNA removal (bacteria) kit from Illumina, and 100-bp single-end read libraries were prepared using a TruSeq Stranded Total RNA Kit (Illumina). The libraries were sequenced using a NextSeq550 sequencer (Illumina).

### Immunolabeling assays

For *Bacillus* immunolabeling assays, poly-L-lysine-coated slides were used. Slides were incubated with poly-L-lysine for 1 h at room temperature and allowed to dry completely for 10 min. Cells were incubated for 24 h at 37 °C with Tse1 and resuspended in PBS. After incubation for 1 h, the cells were fixed in 3% paraformaldehyde and 0.1% glutaraldehyde for 10 min. After two washes in PBS, the samples were then blocked with 3% bovine serum albumin (BSA) and permeabilized with 0.2% Triton-X 100 for 1 h. The slides were then incubated with primary anti-His antibody (rabbit, 1:100 in blocking buffer) for 1 h. Next, the slides were washed three times with washing buffer (0.2% BSA and 0.05% Triton-X 100) and incubated with YFP-fluorescence-conjugated secondary antibody (anti-rabbit, 1:200 in blocking buffer). Finally, cells were stained prior to CLMS with Hoechst 33342 (1:1000) and 100 µg/mL WGA Alexa Fluor 647 conjugate (ThermoFisher), a protein solution that binds to *N*-acetylglucosamine residues in peptidoglycan. For optimal staining, *Bacillu*s cells were washed three times in 3 M KCl before incubation with WGA Alexa Fluor 647 conjugate for 30 min. For immunolabeling assays of Bsub pDR183 + RsiW-YFP, the same protocol was used, but instead using anti-YFP antibody (rabbit, 1:50 in blocking buffer) and YFP-488-conjugated secondary antibody (anti-rabbit, 1:200 in blocking buffer). Image processing was performed using FIJI/ImageJ software (Schindelin *et al*., 2012).

### Statistical analysis

Results are expressed as mean ± standard error of the mean (SEM). Statistical significance was assessed using ANOVA or Student t tests. All analyses were performed using GraphPad Prism® version 9 or Microsoft Excel. P-values < 0.05 were considered significant. Asterisks indicate the level of statistical significance: *p < 0.05, **p < 0.01, ***p < 0.001, and ****p < 0.0001.

## Data availability

All RNA-seq raw data have been submitted to Gene Expression Omnibus (GEO) and can be accessed through GEO series accession no. GSE192348 (URL: https://www.ncbi.nlm.nih.gov/geo/query/acc.cgi?acc=GSE192348).

## Acknowledgements

We thank Saray Morales Rojas for technical support, John Pearson from the Nanoimaging Unit of Bionand for his technical support in the confocal microscopy, Josefa Gómez Maldonado from the Ultrasequencing Unit of the SCBI-UMA for RNA sequencing, and Mercedes Martín Rufián and Casimiro Cárdenas García from the Proteomic Unit of the SCAI-UMA for technical suggestions, protein sequencing, and LC-MS analysis. We are grateful to Prof. Francisco M. Cazorla (University of Málaga, UMA) for kindly providing the wild-type strain *Pseudomonas chlororaphis* PCL1606 and Prof. Patricia Bernal (University of Sevilla, US) for sharing *Pseudomonas putida* KT2440 and *P. putida* KT2440 ΔT6SS and helpful discussion and suggestions. This work was supported by grants from an ERC Starting Grant (BacBio 637971), Plan Nacional de I+D+i of the Ministerio de Ciencia e Innovación (PID2019-107724GB-I00), and Junta de Andalucía (P20_00479). A.I.P.L. is funded by the program FPU (FPU19/00289) and C.M.S. is funded by the program Juan de la Cierva-Incorporación (IJC2018-036923-I).

## Author contributions

D.R. conceived the study; D.R., A.I.P.L. and C.M.S. designed the experiments; A.I.P.L. performed the main experimental work; A.I.P.L and C.M.S. constructed *Bacillus* and *Pseudomonas* strains; D.R., A.I.P.L. and C.M.S. wrote the manuscript; and A.V. contributed critically to writing the final version of the manuscript.

## Expanded view Figure legends

**Table EV1.** Strains used in this study.

**Table EV2.** Oligonucleotides used in this study.

**Dataset EV1.** Upregulated genes after treatment with Tse1.

**Dataset EV2.** Downregulated genes after treatment with Tse1.

**Figure EV1. A)** Schematic representation of T6SS structure. **B)** Representations of T6SS structure during extended, contraction, and disassembly states. The baseplate components (TssA, TssE, TssF, TssG, and TssK) are coloured in blue, the membrane complex (TssJ, TssM, and TssL) in pink and light purple, Hcp monomers in purple, the sheath components (TssB and TssC) in light blue, the spike (VgrG and PAAR) in green, and ClpV in strong purple.

**Figure EV2. A)** Percentages of Bsub sporulation in competition assays using different *Pseudomonas* strains as attackers (*P. chlororaphis* or *P. putida*) demonstrated that *P. putida* strains failed to induce Bsub sporulation **B)** Measurements of the fluorescence signal from *E. coli* pRL662-gfp after incubation with *P. putida* with (KT2440) or without (KT2440 ΔT6SS) a functional T6SS. KT2440 retained toxicity against *E. coli* while KT2440 ΔT6SS failed to kill *E. coli*. **C)** Measurements of the growth of Tse1-expressing *E. coli* (empty square) showed a decrease in cell viability compared with the growth of *E. coli* expressing a truncated version of Tse1 lacking the N-terminus (black square). **D)** Gel electrophoresis of purified His-tagged Tse1 stained with Coomassie and a western blot of purified His-tagged Tse1 exposed to anti-His antibodies (1:1000). Experiments have been repeated at least three times with similar results. Gel images have been cropped and spliced for illustrative purposes. Statistical significance was assessed via t-tests. *p value < 0.1, **p value < 0.01, ***p value < 0.001.

**Figure EV3.** Transmission electron micrographs of Tse1-treated Bsub cells (bottom panels) showed cell wall protuberances and membrane invaginations compared with untreated cells (top panels). Scales are shown in the images.

**Figure EV4. A)** Cytological alterations of Bsub cells after Tse1 treatment. CLSM of untreated (left panel) or Tse1-treated (right panels) *Bacillus* cells stained with Hoechst 33342 (Hoechst, blue channel) (DNA), WGA conjugated with Alexa Fluor 647 (WGA-647, pink channel) (peptidoglycan), or immunolabeling with anti-His antibody conjugated to Alexa Fluor 488 (Anti-His-488, yellow channel) (Tse1 detection). Scale bars equal 20 μm. **B)** Colocalization of Tse1 and peptidoglycan revealed via CLSM and immunodetection of Tse1 and WGA-647 staining. Pearson’s correlation coefficient was calculated to evaluate colocalization between the two channels, Anti-His-488 and WGA-647. Colocalization analysis of the same channel (WGA-647) was used as a positive control for Pearson’s correlation coefficient. Statistical significance was assessed via t-tests. ****p value < 0.0001.

**Fig EV5.** Relative abundance of the peaks found in mass spectrometry analysis of Bsub peptidoglycan treated with buffer (PG, top) or Tse1 (PG + Tse1, bottom). Black arrows designate the 371.23 m/z peak (NAG-NAM-Ala-Glu-mDAP-Ala) and red arrows designate the 415.25 m/z peak (disaccharide tetrapeptide NAG-NAM-Ala-Glu-mDAP). NAG, *N*-acetylglucosamine; NAM, *N*-acetylmuramic acid; Ala, alanine; Glu, glutamic acid; mDAP, *meso*-diaminopimelic acid.

**Figure EV6.** CLSM images of TMRM-stained Bsub cells showing the membrane potential of untreated cells (top) or Tse1-treated Bsub cells (bottom). The data and images in Fig 3E are from the same experiment as the data displayed in this figure and were used to obtain the fluorescence intensity measures. Scale bar equal 20 μm

**Figure EV7. A)** Competition assays between Bsub strains mutant for single kinases (*kinA*, *kinB*, *kinC,* or *kinD*) and Pchl strains (Pchl, Δ*hcp* and Δ*tse1*) showed that single *kinA* and *kinB* mutants are blind to the presence of a functional T6SS or Tse1. **B)** A Coomassie blue-stained SDS denaturing gel showing cell-free supernatant (left) or cell fractions (right) from WT *Bacillus* (Bsub) and *Bacillus* harbouring the RsiW-YFP translational fusion either untreated (Bsub-YFP) or treated with Tse1 (Bsub-YFP + Tse1). Experiments have been repeated at least three times with similar results. Gel images have been cropped and spliced for illustrative purposes. Statistical significance was assessed via t-tests. *p value < 0.1, ***p value < 0.001.

**Fig EV8. A)** Percentages of *Bacillus cereus* (BcerAH) sporulation in competition assays with different Pchl strains as attackers (Pchl, Δ*hcp* and Δ*tse1*). **B-D)** Cell density (CFUs/mL) of different *Bacillus* strains (Bsub, BcerAH, and BcerDSM) in competition assays with Pchl strains (Pchl, Δ*hcp* and Δ*tse1*). Statistical significance was assessed via t-tests. ***p value < 0.001.

